# tRNA m^1^G9 modification depends on substrate-specific RNA conformational changes induced by the methyltransferase Trm10

**DOI:** 10.1101/2023.02.01.526536

**Authors:** Sarah E. Strassler, Isobel E. Bowles, Aiswarya Krishnamohan, Hyejeong Kim, Catherine B. Edgington, Emily G. Kuiper, Clio J. Hancock, Lindsay R. Comstock, Jane E. Jackman, Graeme L. Conn

**Author notes:** For correspondence: Graeme L. Conn,; Jane E. Jackman,.

## Abstract

The methyltransferase Trm10 modifies a subset of tRNAs on the base N1 position of the 9th nucleotide in the tRNA core. Trm10 is conserved throughout Eukarya and Archaea, and mutations in the human gene (*TRMT10A*) have been linked to neurological disorders such as microcephaly and intellectual disability, as well as defects in glucose metabolism. Of the 26 tRNAs in yeast with guanosine at position 9, only 14 are substrates for Trm10. However, no common sequence or other posttranscriptional modifications have been identified among these substrates, suggesting the presence of some other tRNA feature(s) which allow Trm10 to distinguish substrate from nonsubstrate tRNAs. Here, we show that substrate recognition by *Saccharomyces cerevisiae* Trm10 is dependent on both intrinsic tRNA flexibility and the ability of the enzyme to induce specific tRNA conformational changes upon binding. Using the sensitive RNA structure-probing method SHAPE, conformational changes upon binding to Trm10 in tRNA substrates, but not nonsubstrates, were identified and mapped onto a model of Trm10-bound tRNA. These changes may play an important role in substrate recognition by allowing Trm10 to gain access to the target nucleotide. Our results highlight a novel mechanism of substrate recognition by a conserved tRNA modifying enzyme. Further, these studies reveal a strategy for substrate recognition that may be broadly employed by tRNA-modifying enzymes which must distinguish between structurally similar tRNA species.

## INTRODUCTION

RNA modifications are central to proper RNA function and are highly conserved across all kingdoms of life (1). Of all major RNA classes, tRNAs are the most highly modified with 10-20% of tRNA nucleotides containing a modification (2-4). These modifications are critical in determining tRNA fate and tight regulation is crucial for proper cell function due to their roles in ensuring correct codon-anticodon interaction (5,6), fine-tuning tRNA structure and stability (7,8), and regulating tRNA charging with cognate aminoacyl groups (9,10). Because of the abundance of possible tRNA modifications and variations in modification patterns, the mechanisms employed by tRNA modification enzymes to identify their correct tRNA substrates from the large pool of structurally similar tRNAs are still being uncovered.

The Trm10 family of tRNA methyltransferases modify the N1 position of purine nucleotides at position 9 in the core region of tRNA (11). This enzyme family is evolutionarily conserved throughout Eukarya and Archaea, with the first Trm10 enzyme being discovered in *Saccharomyces cerevisiae* (11). Humans express three Trm10 enzymes, with the direct homolog of *S. cerevisiae* Trm10 referred to as TRMT10A and two additional enzymes, TRMT10B and TRMT10C that are distinct in their cellular localization and pool of tRNA substrates. While Trm10/TRMT10A and TRMT10B are both believed to be nuclear/ cytosolic, TRMT10C is localized to the mitochondria as part of the mitochondrial RNase P complex and is the only member of the Trm10 family which is known to function as part of a larger complex (12). Each human Trm10 enzyme also methylates a unique subset of tRNAs, modifying only G9 (Trm10/TRMT10A), only A9 (TRMT10B), or exhibiting bifunctional activity to modify either G9 or A9 (TMRT10C) (13-15).

The importance of Trm10/TRMT10A has been highlighted by its connection to distinct disease phenotypes. In humans, mutations in the *TRMT10A* gene are linked to microcephaly and intellectual disability, as well as defects in glucose metabolism (16-20). Additionally, the *S. cerevisiae trm10* deletion strain exhibits hypersensitivity to the anti-tumor drug 5-fluorouracil (21). While the mechanistic basis of these phenotypes is not fully defined, loss of the m^1^G9 modification increases tRNA fragmentation, which is consistent with the role of core modifications in maintaining optimal tRNA stability (22).

Trm10 is a member of the SPOUT family of *S*-adenosyl-L-methionine (SAM)-dependent methyltransferases which are characterized by an α/β fold with a deep topological knot (23-25). Many SPOUT methyltransferases are involved in RNA posttranscriptional modification (25), including the well-characterized methyltransferase TrmD which modifies G37 of tRNA, catalyzing the same guanosine N1- methylation as Trm10 (11,14). However, TrmD and Trm10 employ very different mechanisms of catalysis. For example, Trm10 does not require a divalent metal ion for catalysis and does not possess the same catalytic residues as demonstrated for TrmD (26-29). Further, in contrast to TrmD and most other SPOUT enzymes, Trm10 is catalytically active as a monomer (30). These differences suggest that Trm10 uses a distinct mechanism of substrate recognition and catalysis.

Trm10 modifies only 14 of 26 possible tRNA substrates in yeast that contain a G9 nucleotide (4,11,14), but no common sequence or posttranscriptional modification(s) have been identified that can explain this substrate specificity (29,30). Trm10 has been shown to efficiently modify *in vitro* transcribed substrate tRNAs, indicating that prior modifications are not necessary for methylation (14). Finally, among tRNAs which do not contain m^1^G9 *in vivo*, there are some which can be modified by Trm10 *in vitro* (hereafter, termed “partial substrates”) and others which are never modified, either *in vitro* or *in vivo* (14). Therefore, some other inherent tRNA property (or properties) must be exploited by Trm10 to discriminate between substrate and nonsubstrate.

Here, we use selective 2’-OH-acylation analyzed by primer extension (SHAPE) to probe inherent tRNA dynamics and their changes upon interaction of Trm10 with substrate and nonsubstrate tRNAs. Our studies demonstrate that there are differences in inherent dynamics in free tRNAs that may allow initial discrimination by Trm10 between substrate and nonsubstrate tRNAs. Using a mutational approach to query the predicted tRNA binding surface of *S. cerevisiae* Trm10, we identify three highly conserved basic residues that are implicated in forming the catalytically productive conformation of the Trm10-tRNA complex. Specifically, the variant protein in which all three residues are altered (Trm10-KRR) is capable of binding tRNAs equivalently to the wild-type enzyme, but fails to significantly modify any substrate tRNA tested, suggesting that formation of the catalytically productive enzyme-substrate complex requires features that were previously unknown. By comparing SHAPE data of tRNA bound to wild-type Trm10 and Trm10-KRR, we show that productive substrate recognition is dependent on the ability of wild-type Trm10 to induce specific tRNA conformational changes to support methylation. These conformational changes are not observed for nonsubstrate tRNA and cannot be induced by the inactive Trm10-KRR variant. Collectively, these studies identify the role of intrinsic and induced tRNA conformational changes and reveal a novel mechanism of RNA substrate recognition by a tRNA-modifying enzyme.

## RESULTS

### SHAPE analysis reveals differences in inherent flexibility of Trm10 substrate and nonsubstrate tRNAs

To assess whether inherent tRNA flexibility might play a role in substrate recognition by Trm10, SHAPE RNA structure probing was performed on five different tRNAs representing substrate (tRNA^Gly-GCC^ and tRNA^Trp-CCA^), nonsubstrate (tRNA^Leu-CAA^ and tRNA^Ser-UGA^), and partial substrate (tRNA^Val-UAC^) (14). These tRNAs were *in vitro* transcribed with short hairpin sequences appended to the 5’ and 3’ ends (**Figure 1A**) to allow measurement of SHAPE reactivities for the full tRNA sequence (31-33). The substrate tRNAs are efficiently modified by Trm10 both *in vivo* and *in vitro* and the nonsubstrate tRNAs are never modified by Trm10 (*in vivo* or *in vitro*). The partial substrate is not modified by Trm10 *in vivo* but modification has been observed *in vitro* with similar reaction kinetics to substrate tRNAs (**Figure 1B**) (14). Additionally, Trm10 activity was not significantly altered by the addition of hairpins to both ends (**Figure 1B**), which is consistent with previous studies showing that SHAPE structure probing of tRNA with similar appended hairpin structures yields consistent results with the authentic tRNA sequences (33).

**Fig. 1.**
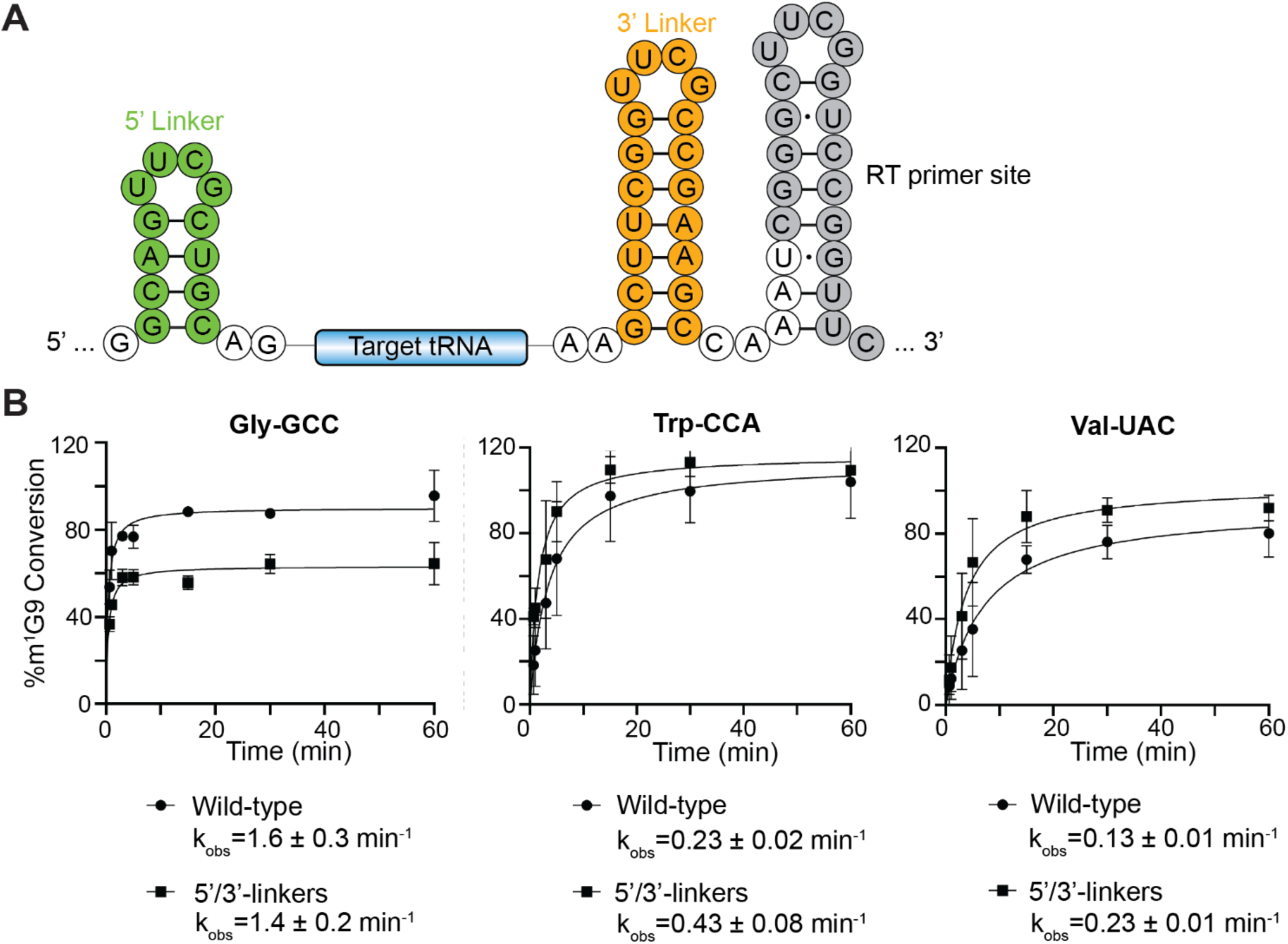
Comparison of modification reaction kinetics for authentic tRNA transcripts and tRNAs embedded within 5’- and 3’-end hairpins. ***A***, Structure and sequence of the *in vitro* transcription template containing 5’- and 3’-hairpins on each side of the tRNA. The RT primer binding site in the 3’-region is also indicated ***B***, Single-turnover reaction plots for authentic tRNA transcripts (wild-type; circles) and SHAPE tRNAs (5’/3’-linkers; squares) for tRNA^Gly-GCC^ (*left*), tRNA^Trp-CCA^ (*middle*), and tRNA^Val-UAC^ (*right*).

SHAPE structure probing was performed on each tRNA using the SHAPE reagent 1-methyl-7- nitroisatoic anhydride (1M7) and analyzed using capillary electrophoresis. The resulting reactivities were determined using RiboCAT software (34) and normalized to the average of the highest reactivities for each sample (**Figure 2A**, and see *Experimental Procedures* for details of the process used). The results were then mapped onto the secondary and tertiary structures of tRNA (**Figure 2B**) and reveal that both substrate and partial substrate tRNAs have high SHAPE reactivity in the 3’-side of the D-loop and the anticodon loop, indicating regions of higher intrinsic tRNA flexibility. These tRNAs also both have low reactivities for nucleotides in the 5’-half of the anticodon stem, core region, acceptor stem, and T-loop indicating more rigid and/ or inaccessible regions. Thus, the intrinsic tRNA dynamics of both substrate and partial substrate tRNAs in the absence of Trm10 appear indistinguishable. In contrast, very different trends for SHAPE reactivity were observed for nonsubstrate tRNAs which exhibit more extensive regions of high SHAPE reactivity for the full D-loop and stem, as well as the core region around the Trm10 target site and parts of the acceptor stem and T-loop.

**Fig. 2.**
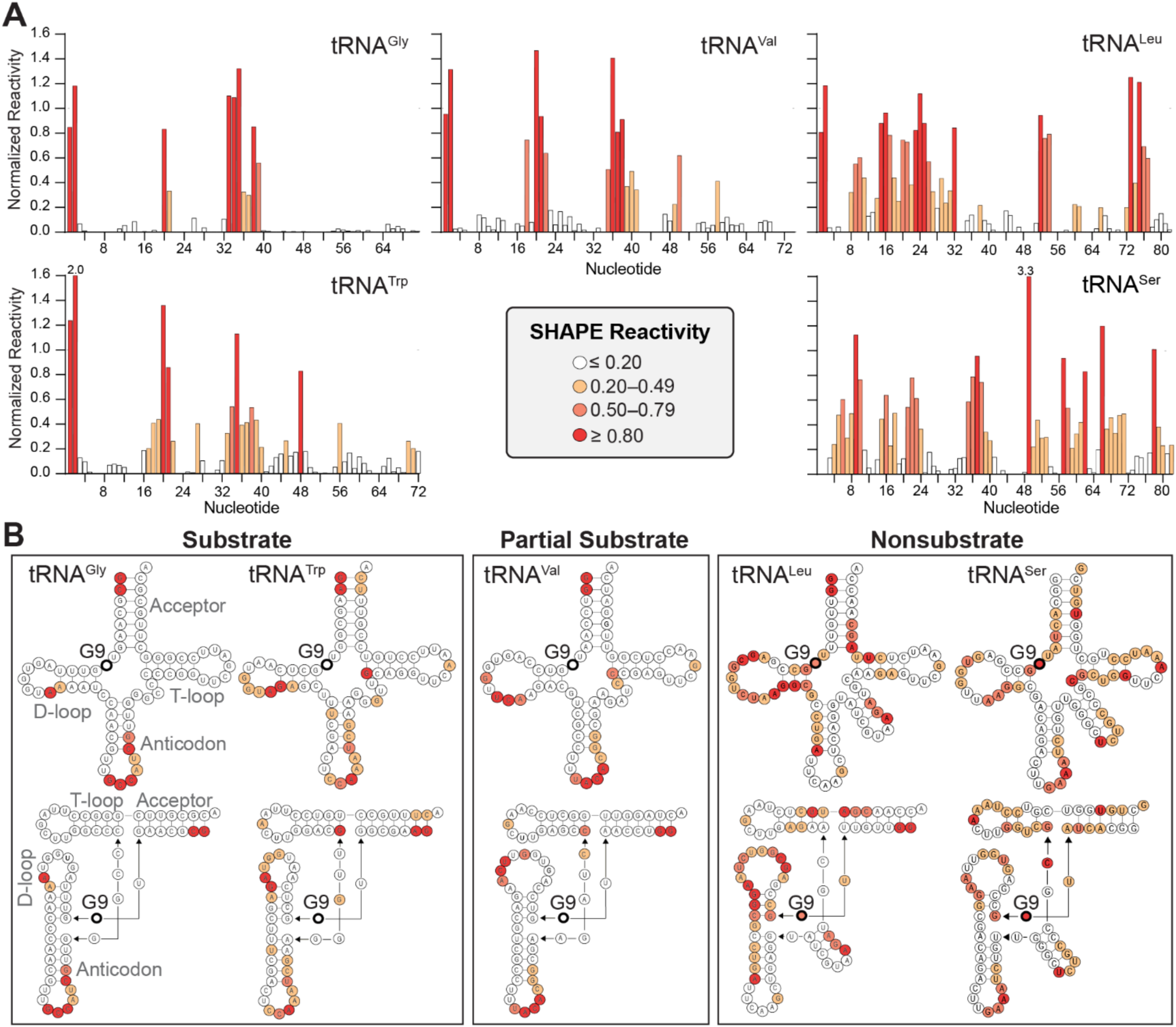
SHAPE analysis reveals differences in inherent flexibility of substrate and nonsubstrate tRNAs. ***A***, SHAPE reactivities for each nucleotide of free tRNA^Gly-GCC^ (substrate), tRNA^Trp-CCA^ (substrate), tRNA^Val-UAC^ (partial substrate), tRNA^Leu-CAA^ (nonsubstrate), and tRNA^Ser-UGA^ (nonsubstrate) were normalized by dividing each value by the average of the reactivities of the highest 8%, omitting the highest 2% (59). The averaged values from two replicates are shown as plots of normalized reactivity vs. nucleotide number. Note that due to the nature of the normalization process, some values may be above 1. The color scale for SHAPE reactivity for both panels is shown in center of the bottom row. ***B***, Averaged SHAPE reactivities for each free tRNA mapped onto their secondary (top) and tertiary (bottom) structures.

These results thus reveal clear differences in inherent tRNA flexibility between substrate and nonsubstrate tRNA which may contribute to substrate discrimination by Trm10. Interestingly, the distinct behavior of the nonsubstrate tRNAs is consistent with the distinct structural nature of nonsubstrate tRNAs, including tRNA^Leu-CAA^ and tRNA^Ser-UGA^, as Type II tRNA species contain an extended variable loop sequence. The observed differences in flexibility for this, and possibly other Type II tRNAs may contribute to the inability of Trm10 to modify any of the tRNAs in this group in any organism studied to date (2,14). The similar inherent flexibility observed with both substrate and partial substrate tRNA is also consistent with the ability of Trm10 to methylate both species *in vitro*. However, these studies with free tRNAs can offer no additional insight into the basis of the substrate preference difference between full and partial substrate *in vivo*. We therefore assessed SHAPE reactivity of the same set of tRNAs in the presence of wild-type Trm10.

### Trm10 induces specific conformational changes in substrate tRNAs that are not observed in nonsubstrate tRNA

To identify changes in tRNA conformation and nucleotide dynamics that occur during Trm10 binding and G9 modification, the same set of five tRNAs (tRNA^Gly-GCC^, tRNA^Trp-CCA^, tRNA^Leu-CAA^, tRNA^Ser-UGA^, and tRNA^Val-UAC^) was pre-incubated with excess Trm10 (at least five times the K_D_ for the protein-RNA interaction; **Supplemental Figure S1**) before adding 1M7 for tRNA SHAPE probing. Additionally, to ensure capture of a homogeneous and catalytically relevant tRNA conformation for the full and partial substrate tRNAs, the SAM cosubstrate-analog “N-mustard 6” (NM6) (35) was included in all SHAPE reactions. *In situ* activation of NM6 results in its covalent attachment to substrates during methyltransferase-catalyzed alkylation reactions, thus trapping the enzyme-substrate-cosubstrate analog complex (35-37) (**Supplemental Figure S2A**). The enzyme-dependent incorporation of NM6 on RNA has been shown previously (37,38) and NM6 was confirmed to be a suitable cosubstrate for tRNA modification by Trm10 using mass spectrometry (**Supplemental Figure S2B**). Thus, NM6 is covalently attached to the tRNA by Trm10 at the N1 base position of G9 such that the Trm10-tRNA complex is trapped in a state immediately following catalysis by virtue of Trm10’s affinity for both the tRNA and the cosubstrate analog which is covalently linked to G9. However, as the SAM analog is only covalently attached to tRNA, Trm10 was removed from the sample after SHAPE modification via phenol:chloroform extraction so that bound protein did not interfere with the subsequent primer extension and capillary electrophoresis analysis. The capillary electrophoresis chromatogram following SHAPE probing shows a large peak at the site of modification (**Supplemental Figure S2C**), providing additional evidence that the SAM analog is incorporated and very efficiently prevents further primer extension (thus no information on reactivity can be determined for subsequent nucleotides 1-8). From these observations we conclude that inclusion of NM6 in SHAPE reactions supports specific capture of an immediate post-catalytic state of the tRNA that accurately reflects changes induced by Trm10.

Analysis and normalization of SHAPE reactivities for each Trm10-bound tRNA was determined as before (**Supplemental Figure S3A** and **S4A,B**) and the corresponding free tRNA values were subtracted. The resulting difference reactivity (Trm10-bound minus free) for each tRNA was then mapped onto the tRNA secondary and tertiary structures (**Figure 3**). Despite being able to bind all tRNAs with similar affinity (**Supplemental Figure S1**), there are significant differences in the conformational changes induced by Trm10 in substrate and nonsubstrate tRNAs. When bound to Trm10, substrates (tRNA^Gly-GCC^ and tRNA^Trp-CCA^) exhibit increased reactivity in the D-loop, particularly in the D-stem immediately adjacent to G9. These changes, which presumably increase accessibility to the target site in the tRNA core, are less pronounced in partial substrate (tRNA^Val-UAC^) and absent in the nonsubstrate tRNAs (tRNA^Leu-CAA^ and tRNA^Ser-UGA^). Additionally, both substrate tRNAs exhibit strong decreases in reactivity in the anticodon stem-loop. This finding was unexpected given the large distance between the anticodon loop and the site of modification, and may be indicative of more global changes in the tRNA structure. This change in the anticodon stem-loop SHAPE reactivity is not observed in partial substrate tRNA^Val-UAC^ or the nonsubstrate tRNAs (tRNA^Leu-CAA^ and tRNA^Ser-UGA^) further suggesting that the corresponding changes in tRNA are important for modification by Trm10. In the former case, we speculate that the observed differences in this region may play a role in this tRNA’s lack of modification *in vivo* where other modifications or interactions may also limit Trm10 activity.

**Fig. 3.**
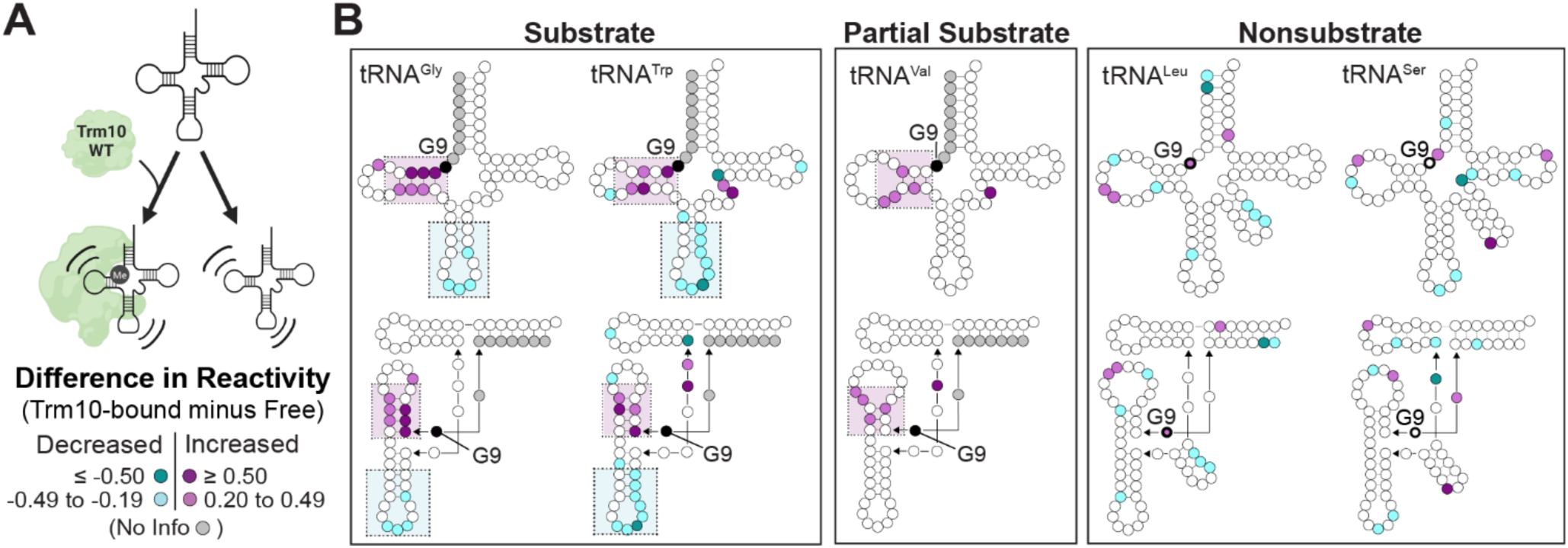
Trm10 induces specific conformational changes in substrate tRNAs that are not observed in nonsubstrate tRNAs. ***A***, Schematic of comparison being made between free tRNA and tRNA bound to wild-type Trm10. ***B***, Difference SHAPE reactivities between Trm10-bound and free tRNAs mapped onto secondary (*top*) and tertiary (*bottom*) structures for (*left* to *right*): tRNA^Gly-GCC^ (substrate), tRNA^Trp-CCA^ (substrate), tRNA^Val-UAC^ (partial substrate), and tRNA^Leu-CAA^ (nonsubstrate), and tRNA^Ser-UGA^ (nonsubstrate). The color scale for difference in SHAPE reactivity (Trm10-bound minus free tRNA) is shown in the *bottom left*.

The SHAPE reactivities may provide some insight into the molecular basis for the observed differences in single turnover rates of modification with the three *in vitro* substrate tRNAs (k_obs_ tRNA^Gly-GCC^ > tRNA^Trp-CCA^/ tRNA^Val-UAC^; **Figure 1B**). We note that for these three tRNAs, there are differences observed in the variable loop which may be important for Trm10 to access and modify G9, as this region is nearby in the folded tRNA structure. Although tRNA^Trp-CCA^ and tRNA^Val-UAC^ exhibit localized increases in SHAPE reactivity, tRNA^Gly-GCC^ exhibits no change in this region upon Trm10 binding. Interestingly, tRNA^Gly-GCC^ has the shortest variable loop (4 nts) of the tRNAs tested, while tRNA^Trp-CCA^ and tRNA^Val-UAC^ are both one nucleotide longer. Thus, the observed changes may reflect a scenario where Trm10 can access G9 of tRNA^Gly-GCC^ without the need to alter its variable loop structure, whereas this region must be altered to allow modification of the other two modified tRNAs (tRNA^Trp-CCA^ and tRNA^Val-UAC^), resulting in the observed differences in activity on these substrate tRNAs. In contrast, with the much longer variable loops of nonsubstrate tRNAs (tRNA^Leu-CAA^ and tRNA^Ser-UGA^), Trm10 is unable to access G9 regardless of its ability to induce conformational changes in this region of the tRNA.

To confirm that the observed differences in SHAPE reactivity between substrate and nonsubstrate are not due to the covalent attachment of NM6 only in substrate tRNA, SHAPE experiments were also performed for a substrate (tRNA^Trp^) and nonsubstrate (tRNA^Leu^) bound to wild-type Trm10 in the presence of the methylation reaction byproduct, S-adenosylhomocysteine (SAH). SAH was selected for this experiment in place of NM6 because it does not result in any modification or covalent attachment of the cosubstrate to the tRNA. The results recapitulate the key observations of the prior experiments for both substrate and nonsubstrate tRNAs, including both increased reactivity in the D-loop and decreased reactivity in the anticodon loop of substrate tRNA^Trp^ (**Supplemental Figure S5**). Interestingly, however, two distinctions are apparent for substrate tRNA with Trm10 in the presence of SAH compared to NM6. First, the reactivity of nucleotide G10 is dramatically reduced. This observation is consistent with major changes in the position of the target nucleotide G9, and thus its interaction with G10, via specific distortions to the tRNA that only occur when G9 is positioned for modification (and, here, trapped in that state by NM6). Second, in the presence of SAH, substrate tRNA^Trp^ shows additional increases in SHAPE reactivity compared to free tRNA and samples with Trm10 and NM6. These additional changes may reflect conformational heterogeneity (e.g. a mixture of free and different bound states) in the presence of SAH, supporting the use of NM6 as a cosubstrate to stabilize a catalytically relevant Trm10-tRNA complex for SHAPE experiments.

Together, these observations suggest that the specific conformational changes observed in both substrate tRNAs, i.e. increased flexibility in the D-loop and decreased anticodon stem-loop flexibility, are essential for substrate recognition by Trm10. Additionally, the variable loop, and Trm10’s ability to induce alterations in its structure where required, may serve as a negative determinant in substrate selection and thus a major reason why tRNA^Leu-CAA^ and other Type II tRNAs are not substrates for Trm10. However, from these data it is not possible to distinguish conformational or nucleotide flexibility changes resulting from Trm10 binding to tRNA vs. those necessary for m^1^G9 methylation.

### Comparison of tRNA bound to Trm10 and Trm10-KRR highlights conformational changes specifically necessary for methylation

Previous structural studies on Trm10 identified a positively charged surface containing conserved residues K110, R121, and R127 as a putative tRNA binding surface (30). Initial binding assays performed with a variant of *Schizosaccharomyces pombe* Trm10 in which all three residues were converted to glutamic acid (K11E/R121E/R127E) indicated that the enzyme completely lost the ability to bind to tRNA, but the impact on catalytic activity was not determined (30). We reconstructed this variant protein in the context of *S. cerevisiae* Trm10 where the same three residues are substituted to glutamic acid (Trm10-KRR; **Figure 4A,B**), and analyzed both tRNA binding and methylation activity. Interestingly, in contrast to the complete loss of tRNA affinity of the *S. pombe* Trm10 variant observed in the previous study, we observed binding of *S. cerevisiae* Trm10-KRR to substrate tRNA^Gly-GCC^ via electromobility shift assay (**Supplemental Figure S6**). While the basis for these distinct impacts of the KRR amino acid changes in the two enzymes is not readily apparent, Trm10 enzymes from *S. pombe* and *S. cerevisiae* only have ∼40% sequence identity and differences in behavior of the “same enzyme” from different organisms have been well documented for other SPOUT methyltransferases (39). The binding of Trm10-KRR to tRNA was further quantified via fluorescence anisotropy-based binding assays which revealed that binding of this Trm10-KRR variant to substrate tRNA^Gly-GCC^ remains essentially identical to that of wild-type Trm10 (**Figure 4C**). However, methylation activity of Trm10-KRR is abolished with tRNA^Gly-GCC^ (**Figure 4D**). Moreover, activity of Trm10-KRR also appears to be decreased significantly compared to wild-type with tRNA^Trp-CCA^ and tRNA^Val-UAC^ tRNAs, based on the loss of the strong stop in primer extension in the SHAPE analyses described below.

**Fig. 4.**
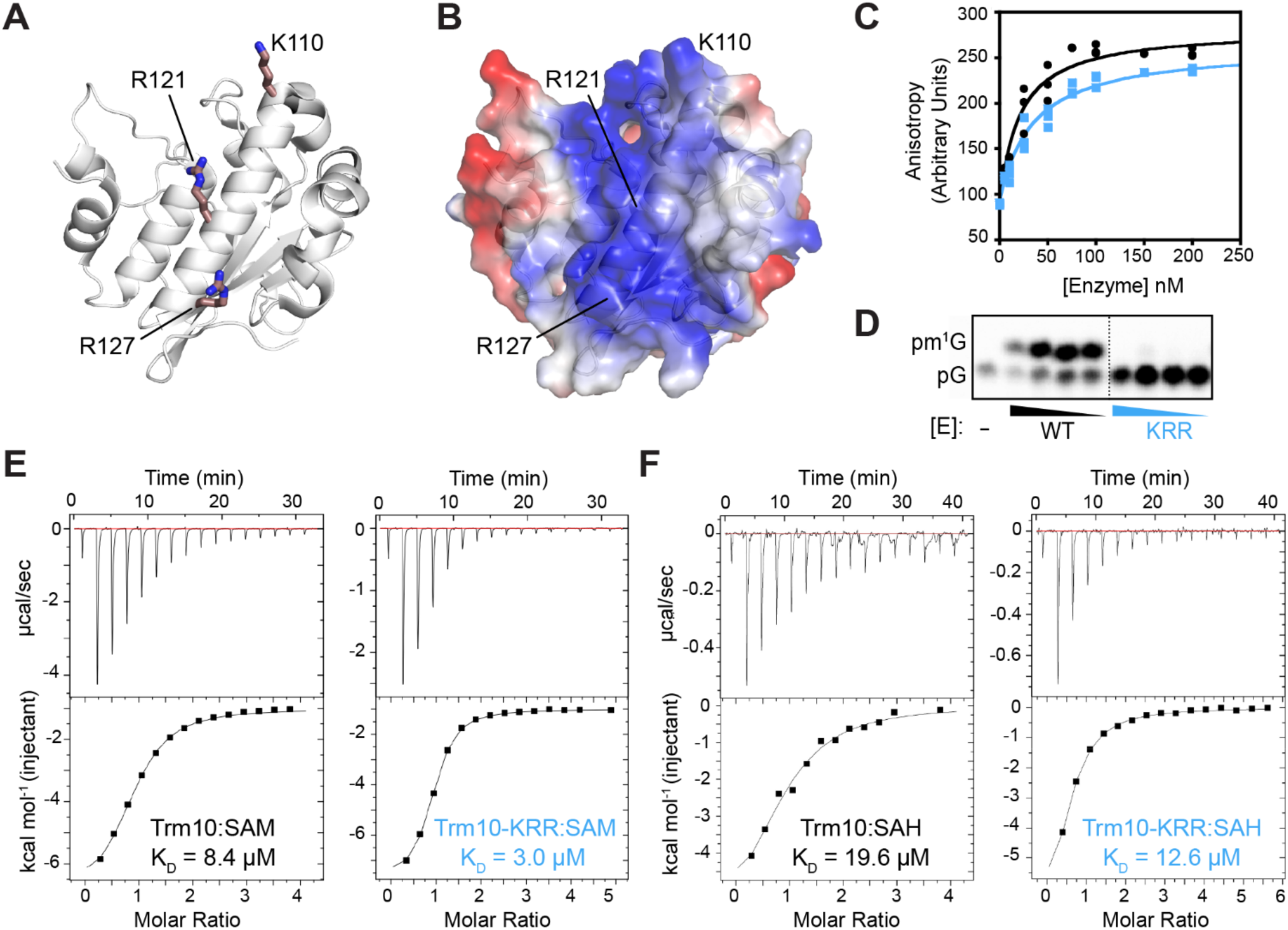
*S. cerevisiae* Trm10-KRR has similar substrate and cosubstrate binding affinities as the wild-type enzyme but lacks catalytic activity. Trm10 C-terminal domain structure (PDB: 4JWJ) shown as ***A***, cartoon and ***B***, electrostatic surface potential highlighting the lysine (K) and arginine (R) residues substituted with glutamic acid in the Trm10-KRR variant. ***C***, Fluorescence anisotropy determination of substrate tRNA^Gly-GCC^ binding affinities (K_D_) for wild-type Trm10 (*black circles*; K_D_ = 20 ± 4 nM) and Trm10- KRR (*blue squares*; K_D_ = 34 ± 5 nM). Results shown are for three independent assays plotted together and fit to equation 1, as described in the Experimental Procedures. ***D***, Thin-layer chromatography methylation assay to determine methylation efficiency of wild-type Trm10 and Trm10-KRR with substrate tRNA^Gly-GCC^. Reactions contained 10-fold serial dilutions of purified enzyme, as indicated by triangles, or no enzyme (-). Representative isothermal titration calorimetry (ITC) analysis of wild-type Trm10 and Trm10-KRR binding to ***E***, SAM cosubstrate and ***F***, methylation reaction by-product SAH. Binding affinities derived from fits to both replicates are given in **Supplemental Table S1**.

Given its comparable tRNA binding affinity to wild-type Trm10, we reasoned that the defect in Trm10-KRR activity could arise either through a defect in enzyme-cosubstrate interaction (e.g. reduced SAM affinity), or an inability to induce changes in the tRNA structure necessary for methylation. To first test the former possibility, we used isothermal titration calorimetry to determine that the affinities of both SAM and SAH are essentially the same (2- to 3-fold differences) for the wild-type Trm10 and Trm10-KRR protein (**Figure 4E,F**). Thus, loss of activity in Trm10-KRR is not due to a defect in SAM/ SAH binding and we propose that the observed impact on enzymatic activity is instead due to an inability to induce some, or all, of the conformational change(s) which are necessary for methylation in these substrate tRNAs after initial Trm10-tRNA binding. As such, Trm10-KRR is a useful probe to dissect changes in tRNA SHAPE reactivity arising from binding and those mechanistically required for modification in tRNA substrates.

To define the tRNA conformational changes specifically necessary for methylation, the same set of tRNAs was pre-incubated with Trm10-KRR before adding 1M7 SHAPE reagent. NM6 was included in these reactions to remain consistent with wild-type Trm10 reactions. Following the same procedures for processing and normalization, SHAPE reactivity of each Trm10-KRR-bound tRNA was mapped onto the tRNA secondary structures (**Supplemental Figure S3** and **S4**). Reactivity differences were calculated for free tRNA and Trm10-KRR bound tRNA (Trm10-KRR-bound minus free; **Supplemental Figure S7**) and both protein-bound states (Trm10-bound minus Trm10-KRR-bound; **Figure 5A,B**). Trm10-KRR appears to induce only a small number of changes in nucleotide flexibility in contrast to wild-type Trm10 (**Supplemental Figure S7**). As such, the calculated differences in tRNA SHAPE reactivities when bound to Trm10-KRR *vs*. Trm10 and Trm10 *vs*. free tRNA are essentially identical (**Figures 3B** and **5B**). Specifically for the protein-bound comparison, the same increases in D-loop and decreases in anticodon stem-loop reactivity are observed. These changes are again absent for nonsubstrate tRNAs and only present in the D-loop of the partial substrate. The variable loop shows an identical trend to the previous comparison for substrate and partial substrate with increased (tRNA^Trp-CCA^ and tRNA^Val-UAC^) or unchanged (tRNA^Gly-GCC^) reactivity in this region. Thus, these results confirm that the conformational changes in the D-loop and anticodon loop observed upon wild-type Trm10 binding to substrate tRNAs are specifically necessary for adoption of a catalytically competent complex for tRNA methylation by Trm10.

**Fig. 5.**
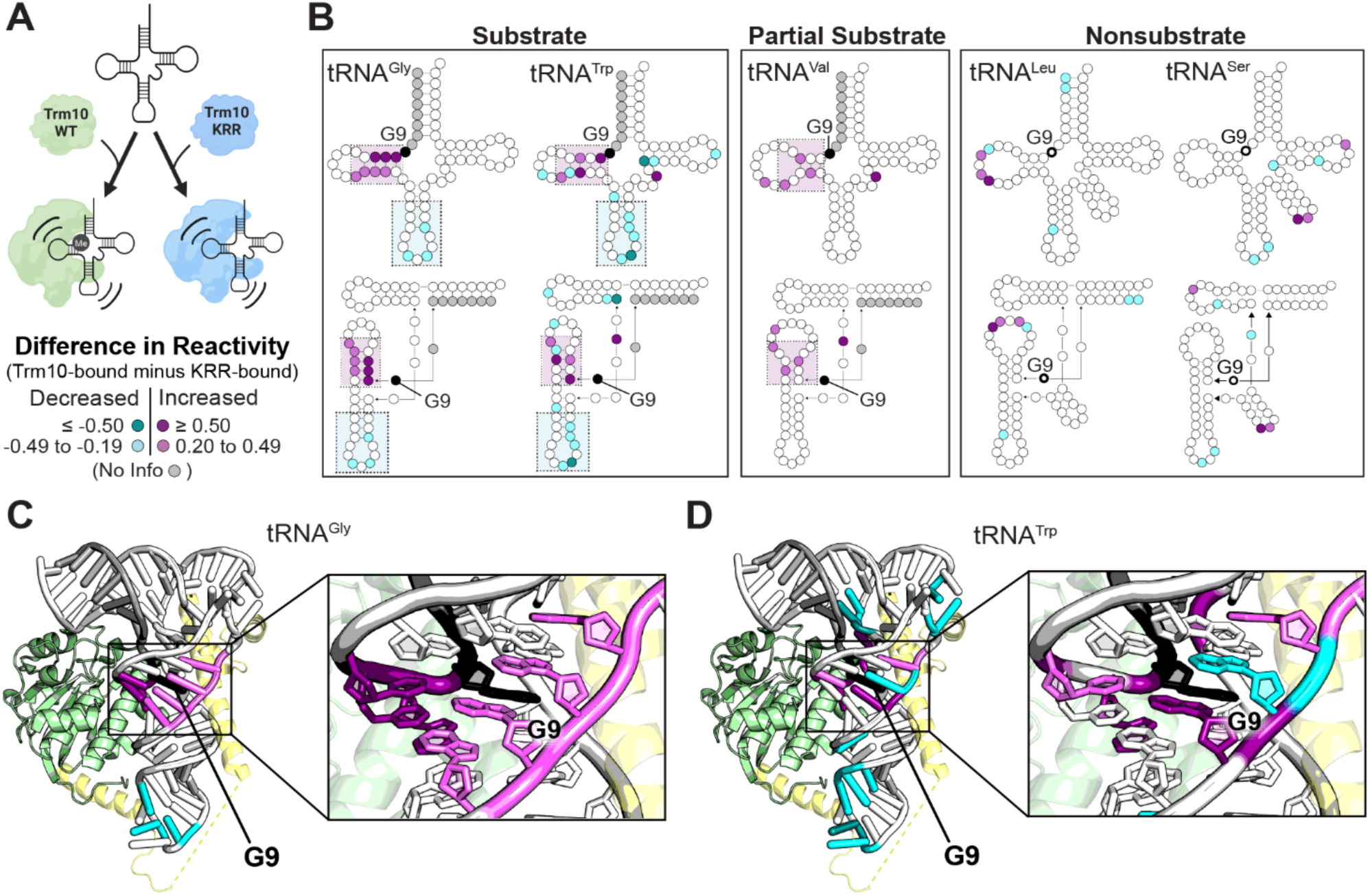
Comparison of SHAPE reactivities when bound to wild-type Trm10 and Trm10-KRR variant reveals the tRNA conformational changes necessary for methylation. ***A***, Schematic of comparison being made between KRR-bound tRNA and Trm10-bound tRNA. ***B***, Difference SHAPE reactivities between Trm10-bound and KRR-bound tRNAs mapped onto secondary (*top*) and tertiary (*bottom*) structures for (*left* to *right*): tRNA^Gly-GCC^ (substrate), tRNA^Trp-CCA^ (substrate), tRNA^Val-UAC^ (partial substrate), tRNA^Leu-CAA^ (nonsubstrate), and tRNA^Ser-UGA^ (nonsubstrate). Difference in reactivity for ***C***, tRNA^Gly-GCC^ and ***D***, tRNA^Trp-CCA^ mapped onto a model of the Trm10-tRNA complex, highlighting conformational changes necessary for methylation. The model of the Trm10-tRNA complex was generated using the structure of TRMT10C bound to pre-tRNA (PDB: 7ONU) and tRNA^Phe^ (PDB: 6LVR), and is comprised of: the Trm10 C-terminal domain (PDB: 4JWJ; *green*), tRNA (*white)* and the N-terminal domain of TRMT10C (*yellow*). The color scale for difference in SHAPE reactivity (Trm10-bound minus KRR-bound tRNA) is shown in the *bottom of panel A*.

### Mapping of SHAPE reactivity onto a Trm10-tRNA model highlights interactions critical for required conformational changes

To further understand the role of tRNA conformational changes in correct substrate recognition and the interactions which support them, a model of Trm10 bound to tRNA was generated using the structure of TRMT10C bound to pre-tRNA as part of the RNase P complex (PDB: 7ONU) (40) (**Figure 5C,D**). This structure was used to guide our modeling as it is currently the only available structure of any Trm10 family member bound to tRNA. The tRNA^Phe^ structure (PDB: 6LVR) was first fit into the binding pocket of TRMT10C by aligning it with the pre-tRNA. Next, the structure of *S. cerevisiae* Trm10 (PDB: 4JWJ) was aligned with TRMT10C to show the placement of this protein with respect to the tRNA. Because the structure of Trm10 lacks the N-terminal domain, the N-terminal domain of TRMT10C was retained in the model to show its approximate location and how the analogous domain of *S. cerevisiae* Trm10 might be positioned to bind the tRNA. Finally, SHAPE reactivity differences corresponding to conformational changes necessary for methylation (i.e. KRR-bound tRNA subtracted from Trm10-bound tRNA reactivity) were mapped onto the structure of the tRNA (**Figure 5C,D**). Although limited due to differences in sequence and the requirement for partner proteins between Trm10 and TRMT10C, this model provides a useful framework for visualizing and interpreting SHAPE reactivity changes observed in the tRNA.

Regions of increased SHAPE reactivity cluster around the site of modification (G9) in the folded tRNA, consistent with an essential role of these Trm10-binding induced conformational changes in allowing accessibility to the target nucleotide. Such changes are also consistent with a mechanism of catalysis that requires the target nucleotide to be rotated 180˚ around its phosphodiester bond (“base-flip") to allow the base to enter the catalytic pocket (41). In further support of this mechanism, we note that the adjacent nucleotide, G10, displays very high reactivity when NM6 is included in the SHAPE reaction (but is absent in the presence of SAH), thus suggesting G9 is secured in the flipped conformation immediately after modification. In our model, the anticodon stem-loop is predicted to interact with the Trm10 N-terminal domain; these interactions may distort the tRNA structure in a way that leads to stabilization or occlusion of this region (decreased SHAPE reactivity) while supporting the changes elsewhere in the tRNA necessary for methylation. Previous studies with a truncated Trm10 protein lacking the N-terminal domain showed a drastic reduction in methylation activity (30), further supporting a model where conformational changes to the anticodon stem-loop induced by the N-terminal domain are essential for catalysis.

## DISCUSSION

One challenge faced by all tRNA-modifying enzymes is how to recognize specific substrates from a pool of tRNAs with a similar overall structure. Sequence and modification recognition elements have been identified for some enzymes, but many key aspects of molecular recognition are still being uncovered (39,42). Trm10 methylates a subset of tRNAs with a guanosine at position 9 and these tRNAs appear to have no common sequence or other posttranscriptional modifications that result in selection of some, but not all, G9-containing tRNAs for modification. Moreover, we previously demonstrated that the long variable loop of Type II tRNAs is a defining feature of nonsubstrate tRNAs (14), but the molecular basis for this distinction had not been demonstrated. Thus, some other common structural feature(s) among substrate tRNAs or among nonsubstrate tRNAs that affect modification, must underpin Trm10’s observed specificity.

Here, we used the SAM-analog NM6 to capture the Trm10-tRNA complex in a catalytically relevant state and showed that methylation by Trm10 is dependent on specific conformational changes to the substrate tRNA that are induced by binding of the enzyme. Our SHAPE structure probing revealed distinct conformational changes in substrate tRNA that are necessary for methylation and which are not observed in Trm10-bound nonsubstrate tRNA. These changes are also not observed for any tRNA in the presence of the tRNA-binding competent but catalytically inactive Trm10-KRR variant. The changes include increased reactivity in the D-loop of the tRNA and decreased reactivity in the anticodon loop, which are consistent with a model in which local conformational changes position the target nucleotide in its binding pocket, while distant conformational changes are related to specific interactions with the N-terminal domain of Trm10.

Considering the inaccessibility of the G9 target nucleotide in the core region of the tRNA, the increased reactivity for nucleotides in the D-loop which surround the site of modification is likely to be necessary for access to the target base and to position it for methylation. Based on the observed nucleotide reactivity changes surrounding G9 in the Trm10-bound substrate tRNA and lack of G10 SHAPE reactivity in the presence of wild-type Trm10 with SAH compared to NM6, we speculate that methylation may require a process known as base-flipping, which is common among DNA and RNA methyltransferases that act on an inaccessible nucleotide. First observed in the methyltransferase *M.HhaI* DNA C5-methyltransferase complexed with a synthetic DNA complex (43), all structures solved since for base-modifying SPOUT methyltransferases in complex with substrate RNAs (TrmD (44), Nep1 (45), and TRMT10C (40)) exhibit this process as part of the modification mechanism. Considering that nonsubstrate tRNAs bound to wild-type Trm10 and substrate tRNAs bound to the inactive Trm10-KRR variant do not show these same increases in reactivity around the site of modification, base-flipping presumably only occurs after initial binding and directly prior to methylation once the correct structural elements have been recognized. The inability of the inactive Trm10-KRR variant to induce the tRNA changes that we propose are necessary to flip the target base into position implies that one or more of the mutated residues is likely essential for driving and/ or recognizing these alterations in the tRNA core region. This idea is supported by comparison of the electrostatic surface potential of wild-type Trm10 and Trm10-KRR (**Supplemental Figure S8**). The observation of tRNA binding to the Trm10 variant suggests that Trm10-KRR maintains enough positive residues around the putative binding surface. Moreover, the replacement of arginine and lysine residues around the catalytic center may prevent the necessary distortion of the tRNA around the site of modification, rendering the target nucleotide inaccessible. As precedence for such a mechanism, basic residues that are critical for RNA distortion, but which do not contribute measurably to RNA substrate binding, have been identified in other RNA-modifying enzymes such as the aminoglycoside-resistance 16S ribosomal RNA methyltransferase RmtC (46,47). However, a precise understanding of the role of these residues in Trm10 requires further detailed investigation.

The decreased reactivity in the anticodon loop distant from the site of modification was more surprising. However, these changes can potentially be explained in the context of our model for Trm10- tRNA interaction (**Figure 5C,D**) which provides a useful framework to visualize tRNA conformational changes and to predict how the tRNA may be interacting with different regions of Trm10. The conformational changes in the anticodon loop may be related to specific interactions made by the N-terminal domain of Trm10 with the opposite surface of the tRNA to that used for C-terminal domain binding. Despite the lower sequence conservation and likely structural differences between the TRMT10C and Trm10 N-terminal domains compared to the C-terminal SPOUT fold, both are enriched in positively charged residues and have been implicated in tRNA binding (30,40). Thus, a reasonable expectation is that the Trm10 N-terminal domain wraps around the anticodon stem-loop to make similar interactions with tRNA as for TRMT10C. Therefore, the region linking the two Trm10 domains may directly interact with the anticodon loop of tRNA, leading to the reduction in reactivity in this region of the tRNA. However, we note that as Trm10 is not known to function as part of larger complexes, there are limitations to the information that can be inferred from this model which was generated using the structure of TRMT10C as part of the larger mitochondrial RNase P complex. Further direct structural information is needed to draw specific conclusions about the interactions between the Trm10 and tRNA. Nonetheless, an essential role for the N-terminal domain in distorting the anticodon loop during recognition would also explain why a truncated Trm10 enzyme which is lacking the N-terminal domain shows a drastic reduction in methylation activity (30). Recognition of the tRNA as a whole and of regions distant from the modification site is common amongst tRNA-modifying SPOUT methyltransferases, including TrmD (48), TrmJ (49,50), and TrmH (51), as well as other tRNA modifying enzymes such as human pseudouridine synthase PUS7 (52). Each of these enzymes require full-length tRNA for efficient modification, implying that they recognize structural elements of the tRNA outside of just the modification site, in a manner similar to Trm10.

It is additionally possible that the interactions between Trm10 and the anticodon loop of substrate tRNA directly contribute to recognition by helping to propagate long distance changes in the tRNA conformation which are necessary for methylation. Conformational changes to the substrate which are distant from the site of modification have been observed for other SPOUT methyltransferases including thiostrepton-resistance methyltransferase, which unfolds the tertiary structure of its substrate ribosomal RNA to cause a more global conformational change (53,54). Similar unfolding processes have also been observed for tRNA-modifying SPOUT methyltransferases, including TrmH (51,55) which modifies the D-loop of tRNA and Trm56 (56) which modifies the T-loop. Both enzymes require disruptions to the tertiary structure of the tRNA for methylation to occur.

Our findings on RNA conformational changes necessary for methylation by Trm10 shed light on a novel component to substrate recognition and can explain why no apparent trends in sequences have been identified to date among substrate or nonsubstrate tRNAs. Very different RNA sequences can result in similar inherent flexibilities in the tRNA structure and/ or capacity to be appropriately reconfigured upon Trm10 binding, as is observed for substrates tRNA^Gly-GCC^ and tRNA^Trp-CCA^. Therefore, consideration of RNA flexibility and deformability as potential recognition elements for a specific RNA-modifying enzyme may be critical in fully defining the recognition process. This may be especially true for other RNA-modifying enzymes which modify an inaccessible region or other tRNA-modifying enzymes which need to be able to discriminate between structurally similar tRNA species.

Thus, while a high-resolution structure is still needed to uncover the details of the interaction between Trm10 and substrate tRNA, the current study has revealed new insights into how Trm10 discriminates between structurally similar tRNAs to select the correct substrate for methylation. Considering the similar overall tertiary structure for all tRNAs, identifying the mechanism of substrate recognition by Trm10 may prove to be critical for understanding substrate recognition for other tRNA-binding proteins as well.

## EXPERIMENTAL PROCEDURES

### RNA in vitro transcription and purification

Genes encoding tRNA^Gly-GCC^, tRNA^Trp-CCA^, tRNA^Val-UAC^, tRNA^Leu-CAA^, and tRNA^Ser-UGA^ were cloned into plasmids for production of *in vitro* transcripts with 5’- and 3’- end hairpins to improve structure probing resolution at the 5’ and 3’ ends of the tRNA sequence (31). All tRNAs were *in vitro* transcribed from *Xho*I linearized plasmid DNA using T7 RNA polymerase as previously described (57). Briefly, *in vitro* transcription was performed for 5 hours at 37˚C in 200 mM HEPES-KOH (pH 7.5) buffer containing 28 mM MgCl_2_, 2 mM spermidine, 40 mM dithiothreitol (DTT), 6 mM each rNTP, and 100 μg/mL DNA template. At the end of the reaction, following addition of EDTA to clear pyrophosphate-magnesium precipitates and dialysis against 1×Tris–EDTA buffer, RNAs were purified by denaturing polyacrylamide gel electrophoresis (50% urea, 1×Tris–Borate–EDTA buffer). RNA bands were identified by UV shadowing, excised, eluted from the gel by crushing and soaking in 0.3 M sodium acetate, and ethanol precipitated as previously described (57).

### Trm10 expression and purification

Trm10 and Trm10-KRR proteins with an N-terminal 6xHis-tag were expressed from the pET-derived plasmid pJEJ12-3 encoding full-length wild-type Trm10 or Trm10-KRR in *E. coli* BL21(DE3) pLysS grown in lysogeny broth as described previously (11). Briefly, protein expression was induced by addition of 1 mM β-D-1-thiogalactopyranoside at mid-log phase growth (OD_600_ ∼0.6) and growth continued at 37˚C for an additional 5 hours. To ensure removal of co-purifying SAM, cells were lysed in buffer containing 1M NaCl and 0.5% TritonX-100, and the lysate dialyzed three times in buffer containing 2M NaCl before dialysis back into original buffer (20 mM HEPES pH 7.5, 0.25 M NaCl, 4 mM MgCl_2_, 1 mM β-mercaptoethanol, 10 mM imidazole, 5% glycerol). Protein was purified by sequential Ni^2+^- affinity (HisTrap HP), heparin-affinity (HiPrep Heparin 16/10), and gel filtration (Superdex 75 16/600) chromatographies on an ÄKTApurifier10 system (GE Healthcare). Trm10 was eluted from the gel filtration column in 20 mM Tris (pH 7.5) buffer containing 100 mM NaCl, 1 mM MgCl_2_, 5 mM β-mercaptoethanol, and 5% glycerol.

### NM6 preparation

The SAM analog NM6 (5’-(diaminobutyric acid)-N-iodoethyl-5’-deoxyadenosine ammoniumhydrochloride) was prepared as previously described (35) and purified by semi-preparative reverse-phase HPLC. Before use, NM6 was dissolved in protein buffer (20 mM Tris pH 7.5, 100 mM NaCl, 1 mM MgCl_2_, 5 mM β-mercaptoethanol, and 5% glycerol). NM6 was included in SHAPE reactions at a final concentration of 5 μM at the same time as Trm10, prior to incubation at 30˚C. *In situ* activation of NM6 results in a Trm10 cosubstrate that is covalently attached by the enzyme to RNA (**Supplemental Figure S2A**), as shown previously (37,38) and confirmed by mass spectrometry (**Supplemental Figure S2B**).

### tRNA SHAPE analysis

SHAPE RNA probing was carried out following previously described procedures (32). Each tRNA was annealed at 95˚C for 2 minutes and incubated on ice for 2 minutes, and addition of folding buffer (333 mM HEPES pH 8.0, 20 mM MgCl_2_, and 333 mM NaCl) followed by incubation at 30˚C for 20 minutes. For samples containing Trm10 or Trm10-KRR, the protein (final concentration of 750 nM) and SAM-analog NM6 or SAH (final concentration 5 μM) were added after RNA folding (35,37). The complex was incubated at 30˚C for 30 minutes before introducing either the SHAPE reagent 1M7 (75 mM), or DMSO control, for 1.5 minutes at 37˚C (58). Trm10 was removed (where required) via phenol chloroform and RNAs recovered by ethanol precipitation.

Reverse transcription (RT) was carried for each product ([+] and [-] SHAPE reagent and one sequencing reaction) with a fluorescently (VIC) labelled primer corresponding to the sequence of the 3’ end of the SHAPE hairpin (**Figure 1A**). Modified RNA was incubated with labeled DNA primer for 5 minutes at 95˚C and cooled to room temperature. The reverse transcription enzyme mix (10 mM dNTPs, 0.1 M DTT, SuperScript III Reverse Transcriptase, and SuperScript III Reverse Transcriptase buffer) was added to the RNA solution and incubated at 55˚C for 1 hour, followed by inactivation at 70˚C for 15 minutes. The resulting cDNA was ethanol precipitated, washed with 70% ethanol, and analyzed by capillary electrophoresis. For capillary electrophoresis, cDNA pellets were resuspended in formamide and mixed with GeneScan^TM^ 600 LIZ® Size Standard (Applied Biosystems) for intercapillary alignment. Samples were resolved on an Applied Biosystems 3730 DNA Analyzer (Genomics Shared Resource Facility, Ohio State University).

Raw electropherograms were converted to normalized SHAPE reactivity using the RNA capillary-electrophoresis analysis tool (RiboCAT) (34). For samples containing the m^1^G9 modification (wild-type-bound substrate and partial substrate tRNAs), a large peak was observed at the modification site (for example, **Supplemental Figure S2C**), blocking the reverse transcriptase and limiting reactivity information 3’ of the site of modification (nucleotides indicated in grey in relevant figures). The SHAPE reactivities were normalized by dividing each value by the average of the reactivities of the highest 8%, after omitting the highest 2% (59).The values from two replicates were then averaged for each nucleotide and the resulting reactivities classified as ≤0.20, 0.20–0.49, 0.50–0.79, and ≥0.80. To compare unbound tRNA, Trm10- bound tRNA, and KRR-bound tRNA, the scaled, averaged reactivities for each nucleotide were subtracted as indicated for each data set. The difference in reactivity for each nucleotide was then classified as decreased (≤ -0.50, or -0.49 to -0.19), no change (-0.20 to 0.19) or increased (0.20 to 0.49, or ≥0.50).

### Isothermal titration calorimetry

Trm10 was dialyzed twice at 4˚C against 20 mM Tris buffer (pH 7.5) containing 150 mM NaCl and concentrated to 50 μM. The final dialysis buffer was used to resuspend SAM and SAH (Sigma-Aldrich) to a final concentration of 1 mM. Experiments were performed on an Auto-iTC_200_ at 25˚C and involved 16 injections of 2.4 μl of SAM or SAH into the cell containing protein. Titrations were performed twice for each combination of protein and ligand. The data were fit to a model for one-binding site in Origin software supplied with the instrument after subtraction of residual heats yielding individual equilibrium association constant (K_a_) values for replicate titrations (reported in **Supplemental Table 1**). A representative titration for each protein-ligand combination is also shown in **Figure 3E,F**.

### Mass Spectrometry (MS)

Samples were prepared for MS by incubating 1 μg of tRNA^Gly-GCC^ with 1.3 μg of wild-type Trm10 alone, or with 100 ng of NM6 prepared in water. Samples were incubated for 1 hour at 37˚C and then digested with 1 unit of RNase T1 (Life Technologies) for 1 hour at 37˚C, and allowed to dry overnight at 40˚C. Samples were resuspended in 0.5% triethylamine in MS grade water before being manually injected for analysis by electrospray MS on a Q Exactive Orbitrap Mass Spectrometer calibrated in negative ion mode. MS spectra were centered on the m/z range corresponding to the masses expected for the unmodified and modified (+393 Da) target site fragment (**Supplemental Figure S3B**).

### Fluorescence anisotropy

tRNA binding was measured by fluorescence anisotropy using wild-type and Trm10-KRR variant proteins and 5’-6-carboxyfluorescein-labeled tRNA^Gly-GCC^ transcripts, prepared, and measured as previously described (17). Reactions containing varied concentration of enzyme (50-300 nM) and 15 nM fluorescently-labeled tRNA were incubated for 30 minutes at room temperature prior to measuring anisotropy using an Infinite M1000 PRO fluorometer (Tecan). Anisotropy was measured and plotted as a function of concentration of each Trm10 enzyme. The data were fit to equation 1 (a modified Hill equation) using Kaleidagraph (Synergy Software) to yield the observed K_D_, the minimum anisotropy (FA_min_), and maximum anisotropy (FA_max_) for each enzyme. The results are plotted from three independent assays performed with each enzyme, with the indicated standard error for the resulting fit.

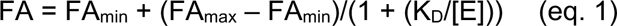

### Electromobility Shift Assay

tRNA^Gly^ was incubated at 80˚C for 10 minutes and then slow cooled to room temperature. The Trm10-tRNA complexes were formed by combining tRNA^Gly^ (4 μM) with final concentrations of wild-type Trm10 or Trm10-KRR ranging from 0 to 10 μM. These mixed components were incubated at 30˚C for 30 minutes to form the complex and then run at 4˚C on a 10% nondenaturing polyacrylamide gel. The gel was incubated with a solution containing ethidium bromide for visualization by UV illumination.

### Trm10 methyltransferase activity assay

Single turnover kinetics were performed as described previously (13,14). Briefly, uniformly labeled transcripts were generated by *in vitro* transcription in the presence of [α-^32^P]-GTP, resulting in labeling of all the G-nucleotides in the tRNA, including at G9. Single turnover reactions (at least 10-fold excess [Enzyme] over [Substrate]) were performed at 30°C with 50 mM Tris pH 8.0, 1.5 mM MgCl_2_, and 0.5 mM SAM. Either 1.5 μM Trm10 (tRNA^Gly-GCC^) or 7 μM Trm10 (tRNA^Trp-CCA^ and tRNA^Val-UAC^) was added to initiate each reaction. At each timepoint, an aliquot of the reaction was quenched by adding to a mixture of excess yeast tRNA and phenol: chloroform: isoamyl alcohol (PCA, 25:24:1 v:v:v). Following PCA extraction and ethanol precipitation, the resulting tRNA was digested to single nucleotides using nuclease P1 (Sigma-Aldrich). Resulting 5’-^32^P labelled p*G or p*m^1^G were resolved by thin layer chromatography on cellulose plates in an isobutyric acid: H_2_O: NH_4_OH solvent (66:33:1, v:v:v). Plates were exposed to a phosphor screen and imaged using a Typhoon imaging system (GE Healthcare). The percent of m^1^G9 conversion at each timepoint was quantified using ImageQuant TL software (GE Healthcare) and the average percent from replicate experiments were plotted versus time using Kaleidagraph (Synergy Software). A k_obs_ value was determined by fitting the plot to a single exponential equation 2, also yielding the P_max_, or maximal amount of product observed for each assay.

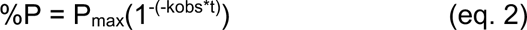

Methyltransferase assays of wild-type Trm10 and Trm10-KRR (**Figure 4D**) were performed essentially identically to the single turnover kinetic assays described above, except that these enzyme titration assays were not performed under enzyme excess conditions and also utilized a tRNA^Gly-GCC^ substrate that was uniquely-labeled with ^32^P at the phosphate immediately 5’ to the G9 nucleotide, as previously prepared and described (27). Reactions were carried out for 2 hours before quenching and processing as described above.

## Data availability

All materials used in this work and the underlying data that support the findings of this study are available from the corresponding author upon request.

## Abbreviations used

1M7: 1-methyl-7-nitroisatoic anhydride
NM6: “N-mustard 6”
MS: mass spectrometry
RT: Reverse transcription
SAH: *S*-adenosylhomocysteine
SAM: *S*-adenosyl-L-methionine
SHAPE: selective 2’-OH-acylation analyzed by primer extension.

## ACKNOWLEDGEMENTS

We thank Drs. Duc Duong and Nick Seyfried for their assistance with MS analysis. This research was supported by the National Institute of General Medical Sciences award R01 GM130135 (to J. E. J. and G. L. C.), the NSF GRFP award 1937971 (to S. E. S.), and the OSU Center for RNA Biology Graduate Fellowship (to I. E. B.). Research reported in this publication was also supported by the Office of the Director, NIH, under award number S10 OD023582.

