## Supporting Information for "tRNA m^1^G9 modification depends on substrate-specific RNA conformational changes induced by the methyltransferase Trm10"

<u>This file contains</u>:

Supplementary Figures S1-S8

Supplementary Table S1

**SUPPLEMENTAL FIGURES**

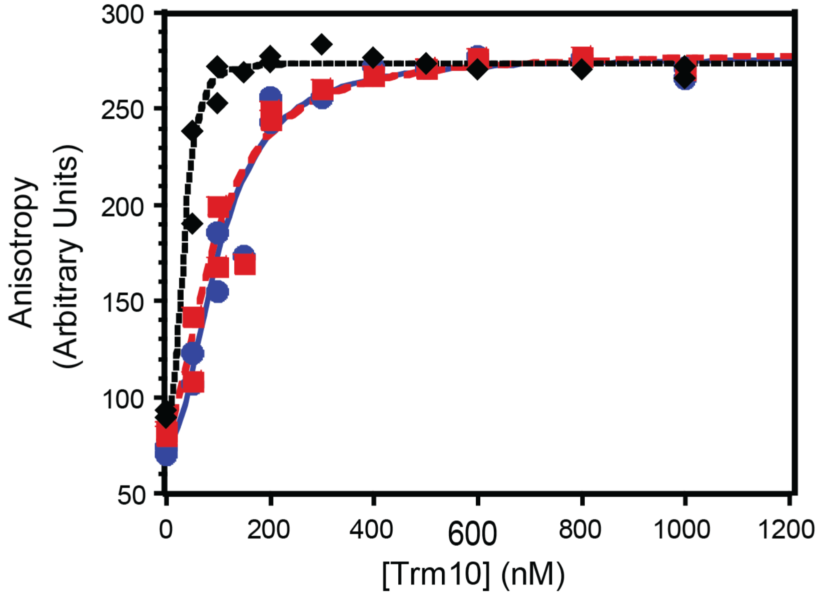


**Supplemental Figure S1. Binding affinity of tRNAs with Trm10.** Wild-type Trm10 binding affinity was determined by fluorescence anisotropy for $tRNA^{Gly\text{-}GCC}$ (*blue circles*; $K_D$ = 100 ± 8 nM), $tRNA^{Val\text{-}UAC}$ (*red squares*; $K_D$ = 96 ± 10 nM), and $tRNA^{Leu\text{-}CAA}$ (*black diamonds*; $K_D$ = 38 ± 5 nM). Data from three independent experiments were plotted together and fit to equation 1 (see *Experimental Procedures*).

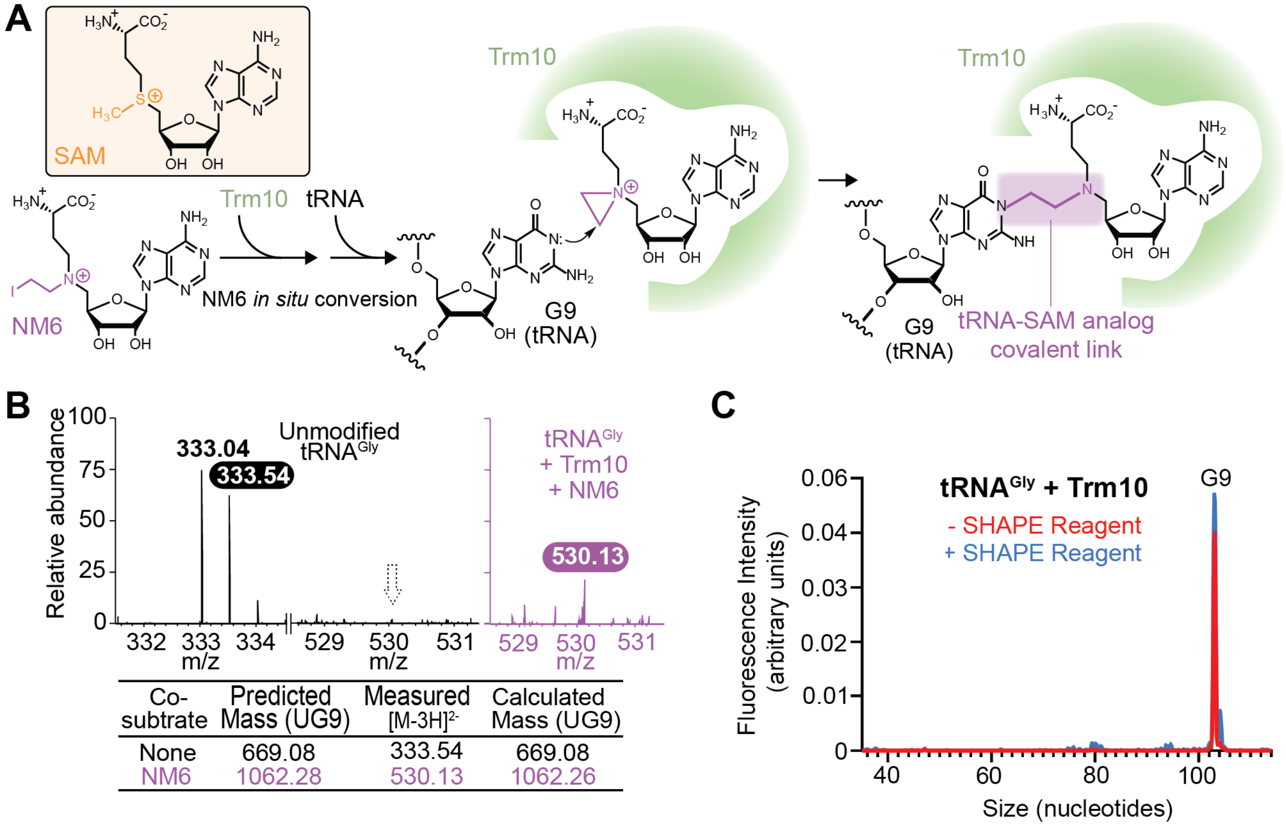


| Co-subtrate | Predicted Mass (UG9) | Measured $[M-3H]^{2-}$ | Calculated Mass (UG9) |
|---|---|---|---|
| None | 669.08 | 333.54 | 669.08 |
| NM6 | 1062.28 | 530.13 | 1062.26 |

**Supplemental Figure S2. Stabilization of the Trm10-tRNA complex using the SAM analog NM6.** ***A***, N-mustard 6 (NM6; *bottom left*) is an analog of S-adenosyl-L-methionine (SAM; *top left box*) that Trm10 can use as a cosubstrate for modification of tRNA. In this enzymatic reaction, NM6 becomes covalently attached to the tRNA at the N1 base position of G9. The Trm10-tRNA complex is thus stabilized in a state immediately following catalysis by virtue of Trm10's affinity for both tRNA subunit and the cosubstrate analog covalently attached to G9. ***B***, The covalent attachment of NM6 to G9 of $tRNA^{Gly}$ is confirmed by MS analysis showing a peak at an m/z of 530.13 representing the $UG_9$-NM6 fragment. This peak is not observed in the sample in which NM6 is absent (dotted arrow). ***C***, Example capillary electrophoresis chromatogram of $tRNA^{Gly\text{-}GCC}$ in the presence of Trm10 and NM6 displaying a large peak at the modification site, indicating that the attachment of NM6 prevents further reverse transcription.

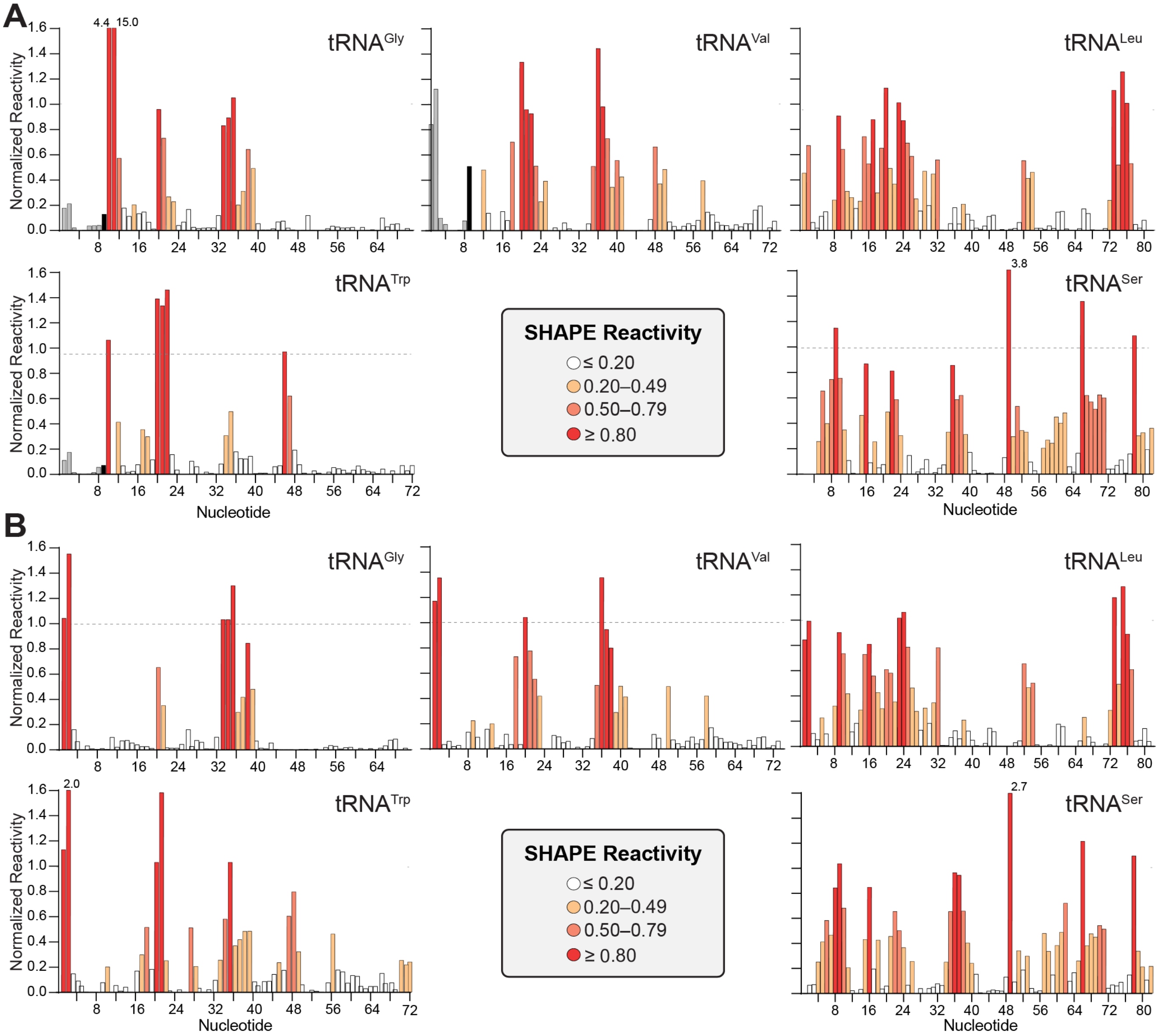


**Supplemental Figure S3. Normalized reactivities of tRNAs bound to wild-type Trm10 and Trm10-KRR.** SHAPE reactivities of tRNA bound to ***A***, wild-type Trm10 or ***B***, Trm10-KRR were normalized by dividing each value by the average of the reactivities of the highest 8%, omitting the highest 2%. The values from two replicates were then averaged and classified as shown in the key in the center of the bottom row of each panel.

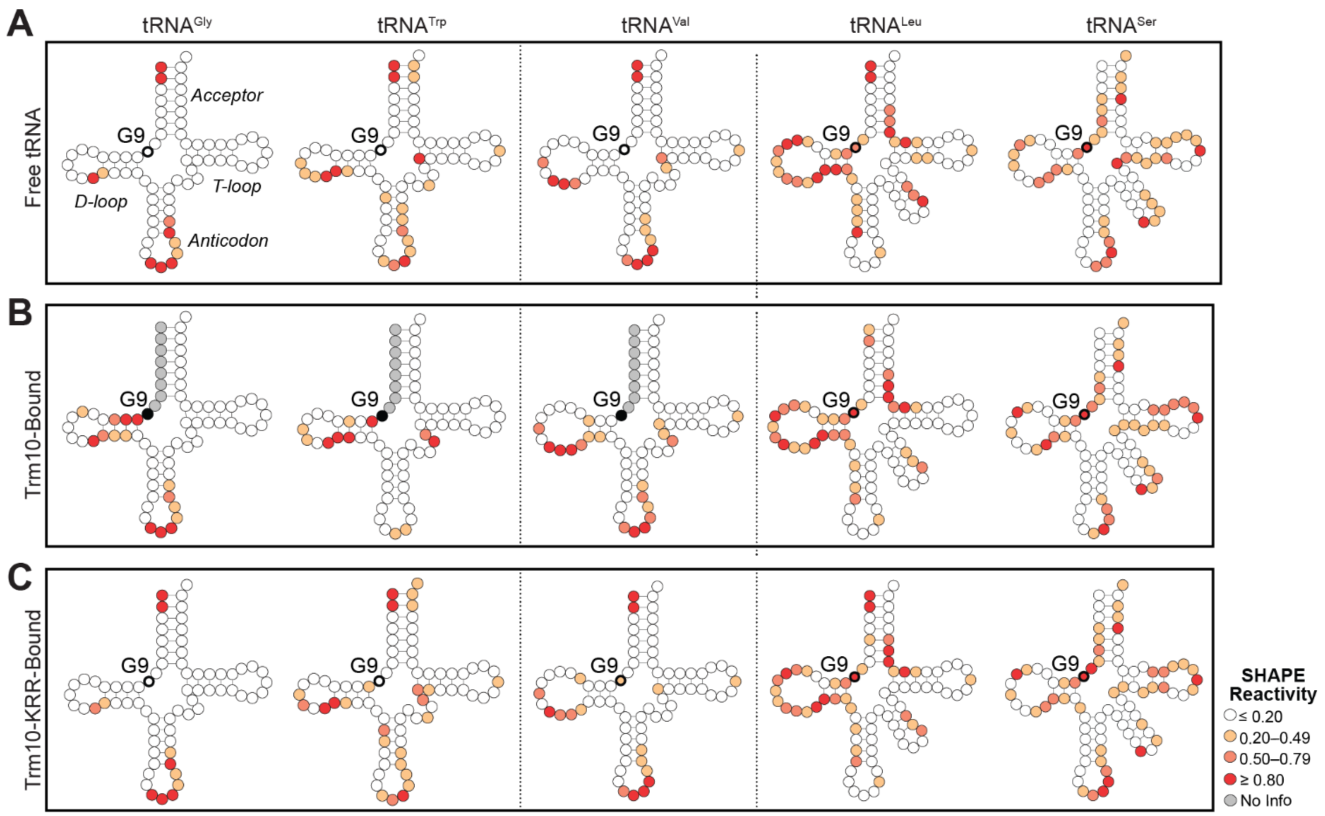


**Supplemental Figure S4. Nucleotide SHAPE reactivities in free and Trm10-bound tRNAs.** SHAPE reactivities mapped onto tRNA secondary structures for: ***A***, free tRNA (note, these data are the same as shown in **Fig. 2B**), ***B***, tRNA bound to wild-type Trm10, and ***C***, tRNA bound to Trm10-KRR. The color scale for SHAPE reactivity is shown in the *bottom right.*

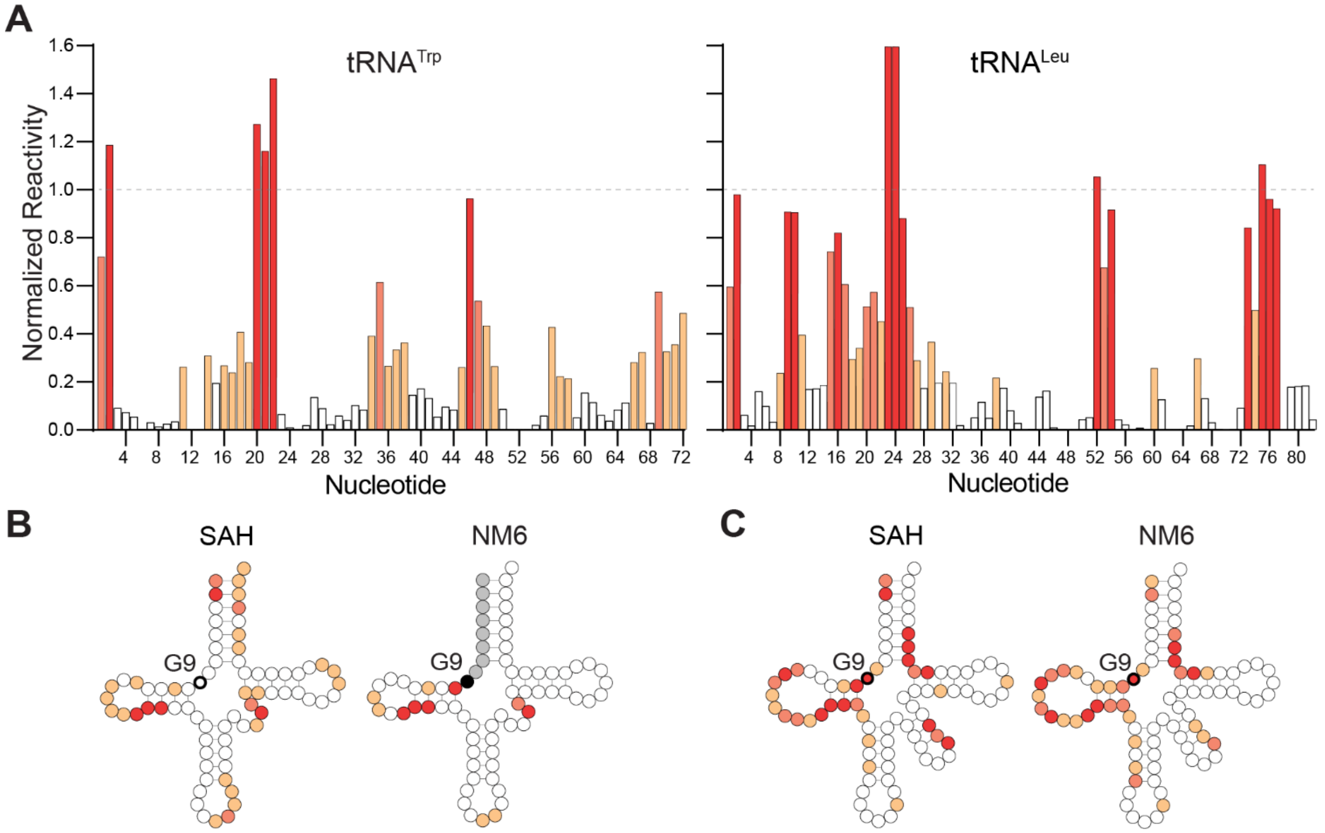


**Supplemental Figure S5. SHAPE reactivities of substrate $tRNA^{Trp}$ and nonsubstrate $tRNA^{Leu}$ bound to wild-type Trm10 in the presence of SAH.** ***A***, SHAPE reactivities of $tRNA^{Trp}$ (substrate) and $tRNA^{Leu}$ (nonsubstrate) bound to wild-type Trm10 in the presence of SAH were normalized and the values for two replicates were averaged. SHAPE reactivities of ***B***, $tRNA^{Trp}$ and ***C***, $tRNA^{Leu}$ bound to wild-type Trm10 in the presence of cosubstrate SAH or NM6 (note, these data for NM6 are the same as shown in **Supplemental Figure S4B**) are mapped onto the tRNA secondary structure. The color scale for SHAPE reactivity is shown for both panels in the *bottom right.*

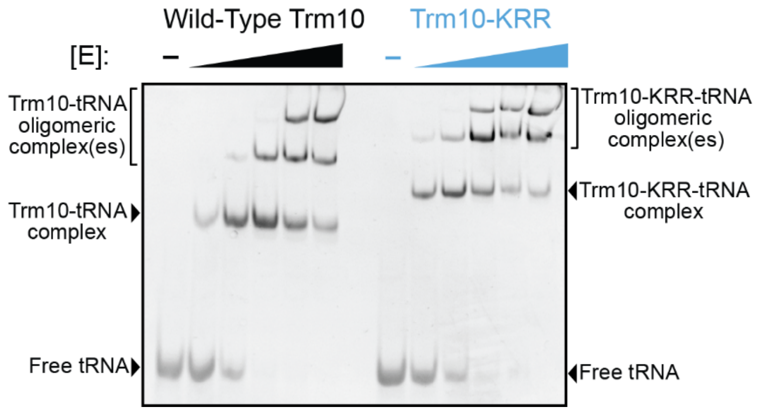


**Supplemental Figure S6. Electromobility shift assay (EMSA) of substrate $tRNA^{Gly}$ with wild-type Trm10 and Trm10-KRR.** $tRNA^{Gly}$ was incubated with increasing concentrations of wild-type Trm10 or Trm10-KRR and samples were run on a non-denaturing 10% polyacrylamide gel. The apparent differences in mobility of the mutant complex are likely due to the different electrostatics of the enzyme-tRNA complex(es) but overall similar binding affinities and patterns of complex formation are observed between wild-type Trm10 and Trm10-KRR.

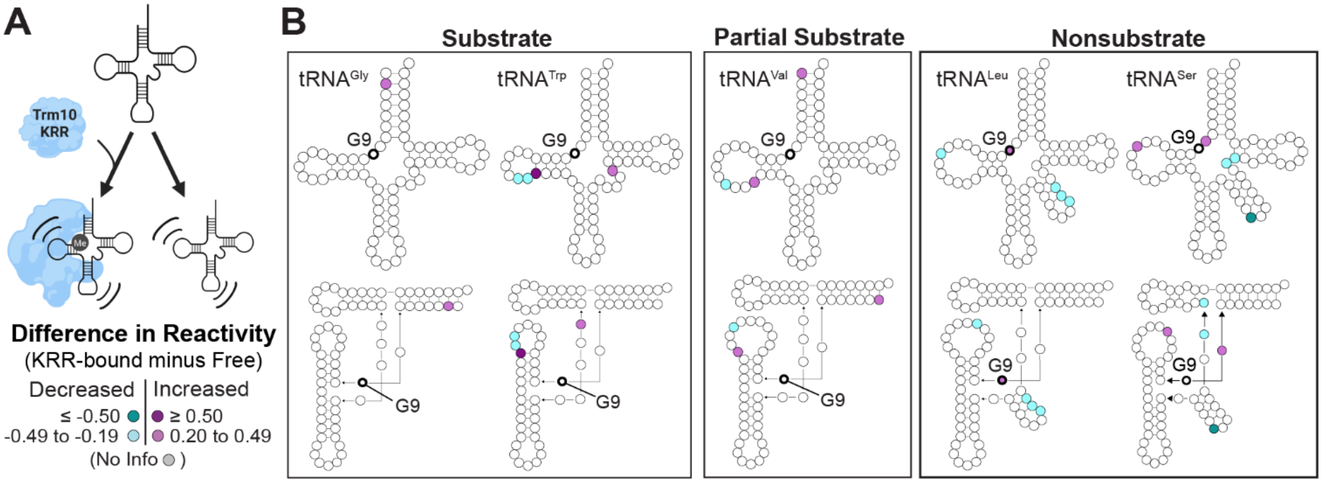


**Supplemental Figure S7. Comparison of unbound tRNA and KRR-bound tRNA.** ***A***, Schematic of comparison being made between free tRNA and KRR-bound tRNA. ***B***, Difference SHAPE reactivities between KRR-bound and free tRNAs mapped onto secondary (*top*) and tertiary (*bottom*) structures for (*left* to *right*): $tRNA^{Gly\text{-}GCC}$ (substrate), $tRNA^{Trp\text{-}CCA}$ (substrate), $tRNA^{Val\text{-}UAC}$ (partial substrate), $tRNA^{Leu\text{-}CAA}$ (nonsubstrate), and $tRNA^{Ser\text{-}UGA}$ (nonsubstrate). The color scale for difference in SHAPE reactivity (KRR-bound minus free tRNA) is shown in the *bottom left*.

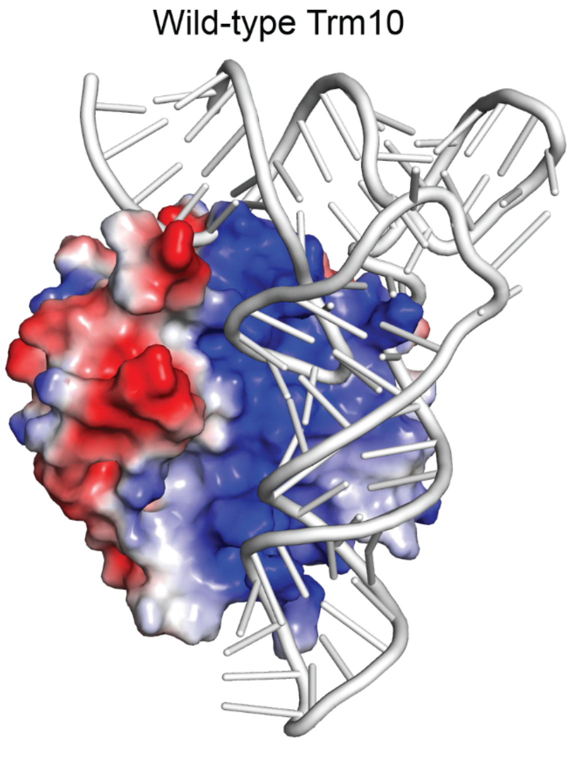


**Supplemental Figure S8. Electrostatic surface potential of wild-type Trm10 and Trm10-KRR shown on a Trm10-tRNA model.** Amino acid substitutions to *S. cerevisiae* Trm10 C-terminal domain (PDB 4JWJ) were made using the PyMol mutagenesis tool to generate the Trm10-KRR variant. Protein electrostatics were then generated for wild-type Trm10 and Trm10-KRR in PyMol and are shown on a semi-transparent surface representation for each protein. The placement of tRNA was predicted from the structure of TRMT10C in complex with pre-tRNA (PDB: 7ONU) and is shown using $tRNA^{Phe}$ (PDB: 6LVR).

### SUPPLEMENTAL TABLES

**Supplemental Table 1. Association constants ($K_A$) for Trm10 binding SAM and SAH determined by ITC.**

| | | SAM[a] | SAH[a] |
|---|---|---|---|
| $K_A$ ($M^{-1}$) | Wild-Type Trm10 | 8.90 ± 0.27 x $10^4$<br>11.9 ± 0.87 x $10^4$ | 5.1 ± 1.42 x $10^4$<br>4.62 ± 1.33 x $10^4$ |
| | Trm10-KRR | 32.5 ± 1.83 x $10^4$<br>33.1 ± 0.88 x $10^4$ | 10.7 ± 1.92 x $10^4$<br>7.96 ± 0.70 x $10^4$ |

[a]Replicate measurements shown with the direct fit error determined using a model for one-binding site in Origin software after subtraction of residual heats.
